# Conserved Roles of Murine SAMHD1 Isoforms in Replication Fork Dynamics and Nascent DNA Processing

**DOI:** 10.64898/2026.11.12.766330

**Authors:** Bishal Manandhar, Domenico Rizzo, Arezoo Jamali, Nina V. Fuchs, Maike Herrmann, Kerstin Schott, Florian D. Hastert, Qun Li, Krisztina Ambrus, Catharina Majer, Jana Kniewel, Thomas Roesler, Alina Henrich, Melanie Rall-Scharpf, Baek Kim, Lisa Wiesmüller, Renate König

**Affiliations:** Host-Pathogen Interactions, Paul-Ehrlich-Institut, 63225 Langen, Germany; Department of Pediatrics, School of Medicine, Emory University, Atlanta 30322 USA; Translational Oncology at University medicine (TRON) of the Johannes-Gutenberg-University, 55131 Mainz, Germany; Department of Obstetrics and Gynecology, Ulm University, 89075 Ulm, Germany; Center for ViroScience and Cure, Children’s Healthcare of Atlanta, 30322 Atlanta, USA

**Author notes:** equal contribution (Domenico Rizzo, Arezoo Jamali, Nina V. Fuchs). Address correspondence to Renate König.

**Keywords:** Mouse SAMHD1, glioblastoma cell model, DNA replication, DNA end resection, isoforms

## Abstract

Sterile alpha motif and histidine–aspartic acid domain-containing protein 1 (SAMHD1) is a deoxynucleoside triphosphate triphosphohydrolase (dNTPase) involved in intracellular nucleotide homeostasis, antiviral restriction, and genome maintenance. While human SAMHD1 (hSAMHD1) contributes to DNA replication fork stability and DNA double-strand break repair, the corresponding functions of murine SAMHD1 (mSAMHD1) remain poorly characterized. Here, we generated SAMHD1-deficient murine GL261 glioma cells using CRISPR/Cas9 to investigate mSAMHD1 function in DNA replication. SAMHD1 knockout increased intracellular dATP levels, altered replication fork dynamics, and reduced camptothecin (CPT)-induced nascent DNA degradation, supporting conserved replication-associated functions. RNA sequencing detected transcripts corresponding to all three annotated *mSamhd1* isoforms, with isoform 1 predominating, followed by isoforms 2 and 3. To assess isoform-specific functions, individual murine isoforms were expressed in *SAMHD1*-deficient human HEK293T cells, enabling direct comparison in a *SAMHD1*-null background. Both major murine isoforms, mSAMHD1 isoforms 1 and 2, localized to the nucleus, exhibited nucleotide-dependent oligomerization and retained dNTPase activity. Notably, both isoforms restored replication fork dynamics and camptothecin (CPT)-induced nascent DNA degradation to levels largely comparable to hSAMHD1. In contrast, isoform 3 exhibited markedly reduced ectopic expression, consistent with its low endogenous transcript abundance in primary murine cells, limiting functional characterization. Together, these findings demonstrate that the two major murine SAMHD1 isoforms retain replication-associated functions of hSAMHD1 despite differences in their C-terminal regions. Furthermore, the established SAMHD1-deficient GL261 cells provide a model for future studies of SAMHD1 function in glioma and its tumor immune microenvironment.

**Importance:** Although SAMHD1 has been extensively studied in human cells, its functions in murine DNA replication and replication fork maintenance remain incompletely understood. Here, we demonstrate that SAMHD1 deficiency impairs replication fork dynamics and reduces stress-induced nascent DNA degradation in murine glioma cells. We further characterize the expression of three annotated murine SAMHD1 isoforms and show that the two major isoforms can restore replication-associated functions in SAMHD1-deficient human cells. These findings support the functional conservation of SAMHD1 between humans and mice and establish a murine cellular model for investigating its contribution to genome maintenance. The generated GL261 SAMHD1-knockout cells provide a foundation for future studies of SAMHD1 in immunocompetent glioma models, where its effects on tumor biology and immune regulation can be investigated.

## Introduction

In humans, the 626-amino-acid sterile alpha motif (SAM) and histidine–aspartic acid (HD) domain-containing protein 1 (SAMHD1) is a conserved deoxynucleoside triphosphate triphosphohydrolase (dNTPase) with important functions in nucleotide homeostasis and antiviral restriction (1, 2). SAMHD1 restricts human immunodeficiency virus type 1 (HIV-1) by depleting intracellular dNTP pools (1, 3, 4), although dNTPase activity alone is not sufficient to confer full antiviral activity (5–7). In addition, phosphorylation of the C-terminal threonine residue T592 has been linked to regulation of HIV-1 restriction (5, 8).

Beyond antiviral restriction, SAMHD1 contributes to genome maintenance. It promotes DNA end resection and homologous recombination through interaction with CtIP (9)and facilitates processing of stalled replication forks by promoting MRE11-dependent degradation of nascent DNA (10). The latter function limits the accumulation of aberrant nucleic acids that may trigger innate immune responses. SAMHD1 also negatively regulates innate immune signaling, including NF-κB and IRF7 activation (11). Consistent with these functions, germline *SAMHD1* mutations cause the hereditary interferonopathy Aicardi–Goutières syndrome (12), whereas somatic *SAMHD1* mutations have been identified in several human cancers (13), linking SAMHD1 dysfunction to autoimmunity and cancer (13–15).

Compared with hSAMHD1, considerably less is known about the cellular functions of murine SAMHD1 (mSAMHD1). Both proteins share the conserved SAM and HD domains and regulate intracellular dNTP levels. Accordingly, *Samhd1*-deficient mice show elevated dNTP levels and altered susceptibility to retroviral infection (16, 17). These mice also display spontaneous type I interferon (IFN-I) activation and increased expression of interferon-stimulated genes (ISGs) (17, 16). This response was initially shown to depend on cGAS-STING signaling (18), while subsequent work identified MDA5/MAVS as the driver of spontaneous IFN-I production, with cGAS-STING-dependent tonic IFN signaling required for MDA5 priming (19). Thus, key functions of SAMHD1 in nucleotide homeostasis and innate immune regulation are conserved between humans and mice.

Nevertheless, hSAMHD1 and mSAMHD1 differ in several biochemical properties. In the absence of allosteric nucleotides, mSAMHD1 predominantly forms dimers, whereas hSAMHD1 is primarily monomeric (20). Moreover, while the isolated HD domain of hSAMHD1 retains tetramerization and dNTPase activity, removal of the SAM domain from mSAMHD1 strongly impairs both processes (20). Thus, the SAM domain has a more prominent role in the oligomerization and catalytic activity of mSAMHD1.

Evidence for a conserved role in genome maintenance is more limited. *Samhd1*-deficient mice exhibit low-level chronic DNA damage, and *Samhd1*-deficient murine macrophages show signs of replication stress (19). However, whether mSAMHD1 directly regulates DNA replication dynamics and DNA end resection, as described for hSAMHD1, remains unclear.

A further difference between human and murine SAMHD1 concerns alternative splicing. Three mSAMHD1 isoforms are currently annotated in the NCBI database, of which the two larger isoforms (627 and 620 amino acids) have previously been described (21, 3). These isoforms arise from alternative splicing of the terminal coding exon and differ in their C-terminal regions. Both retain the SAM and HD domains and are capable of depleting intracellular dNTP pools (22) (23, 3), although differences in phosphohydrolase activity have been reported (22, 20). Notably, mSAMHD1 isoform 2 lacks T603, corresponding to the regulatory T592 phosphorylation site of hSAMHD1 (23) (22) (24). Its absence may therefore affect regulation of mSAMHD1 isoform 2. However, whether the murine isoforms differ in other cellular functions remains unknown, and expression and function of the third annotated isoform have not yet been characterized.

In this study, we generated SAMHD1-deficient murine GL261 glioma cells using CRISPR/Cas9 to investigate the role of mSAMHD1 in DNA replication dynamics and DNA end resection. We selected GL261 cells as a tumor-relevant model that can also be used in syngeneic, immunocompetent mouse models, providing a foundation for future investigations of SAMHD1 function in glioma. We further quantified the annotated *mSamhd1* isoforms in GL261 and primary murine cells and compared their protein functions with hSAMHD1. To assess isoform-specific effects independently of endogenous SAMHD1, individual mSAMHD1 isoforms were expressed in SAMHD1-deficient human HEK293T cells, providing a *SAMHD1*-null background for direct comparison of their roles in replication fork dynamics and stress-induced nascent DNA degradation. Thus, we investigated whether genome maintenance functions established for hSAMHD1 are conserved in mSAMHD1 and whether these functions differ between murine isoforms.

## Results

### Functional consequences of SAMHD1 knockout in murine GL261 cells

Compared with hSAMHD1, the functions of mSAMHD1 have mainly been characterized with respect to dNTPase activity and innate immune regulation, whereas its role in DNA replication remains largely unexplored. To investigate the role of mSAMHD1 in DNA replication, we first generated SAMHD1-protein knockout (SAMHD1-KO) murine glioblastoma GL261 cells.

Using CRISPR/Cas9, we targeted exon 4 of murine *Samhd1*. The generated clones were screened for loss of SAMHD1 expression by immunoblotting and further characterized by PCR-based TA cloning followed by Sanger sequencing (Supplementary Fig. 1A). Immunoblot analysis identified samples #3, #4, and #5 as lacking detectable SAMHD1 protein, whereas cell samples #1 and #2 retained SAMHD1 expression (Fig. 1A and Supplementary Fig. 1B). Sanger sequencing of the SAMHD1-negative samples revealed heterogeneous genomic alterations, including one-or two-nucleotide insertions resulting in premature stop codons in exon 5 as well as deletions affecting the splice donor site. In addition, deletions within exon 4 resulted in a shortened exon 4 sequence (Supplementary Fig. 1C). The presence of different *Samhd1* alleles indicates that the generated SAMHD1-deficient GL261 cells represent a mixed population rather than a single genetically homogeneous clone. Nevertheless, no detectable SAMHD1 protein expression was observed neither with a monoclonal antibody (Fig. 1A), nor with a polyclonal antibody (Supplementary Fig. 1B) confirming the successful generation of SAMHD1-deficient GL261 cells. To assess the functional consequences of mSAMHD1 deficiency, we first examined intracellular dATP levels in SAMHD1-knockout (KO) GL261 cells. Using a single-nucleotide incorporation assay (25), we quantified dATP in cell lysates. Consistent with previous reports of elevated dNTP levels in *Samhd1*-deficient mice (16, 17), SAMHD1-KO cells exhibited a significant increase in dATP levels compared with mock-edited controls (Fig. 1B). These findings are consistent with impaired SAMHD1-dependent dNTP hydrolysis following knockout.

**Figure 1:**
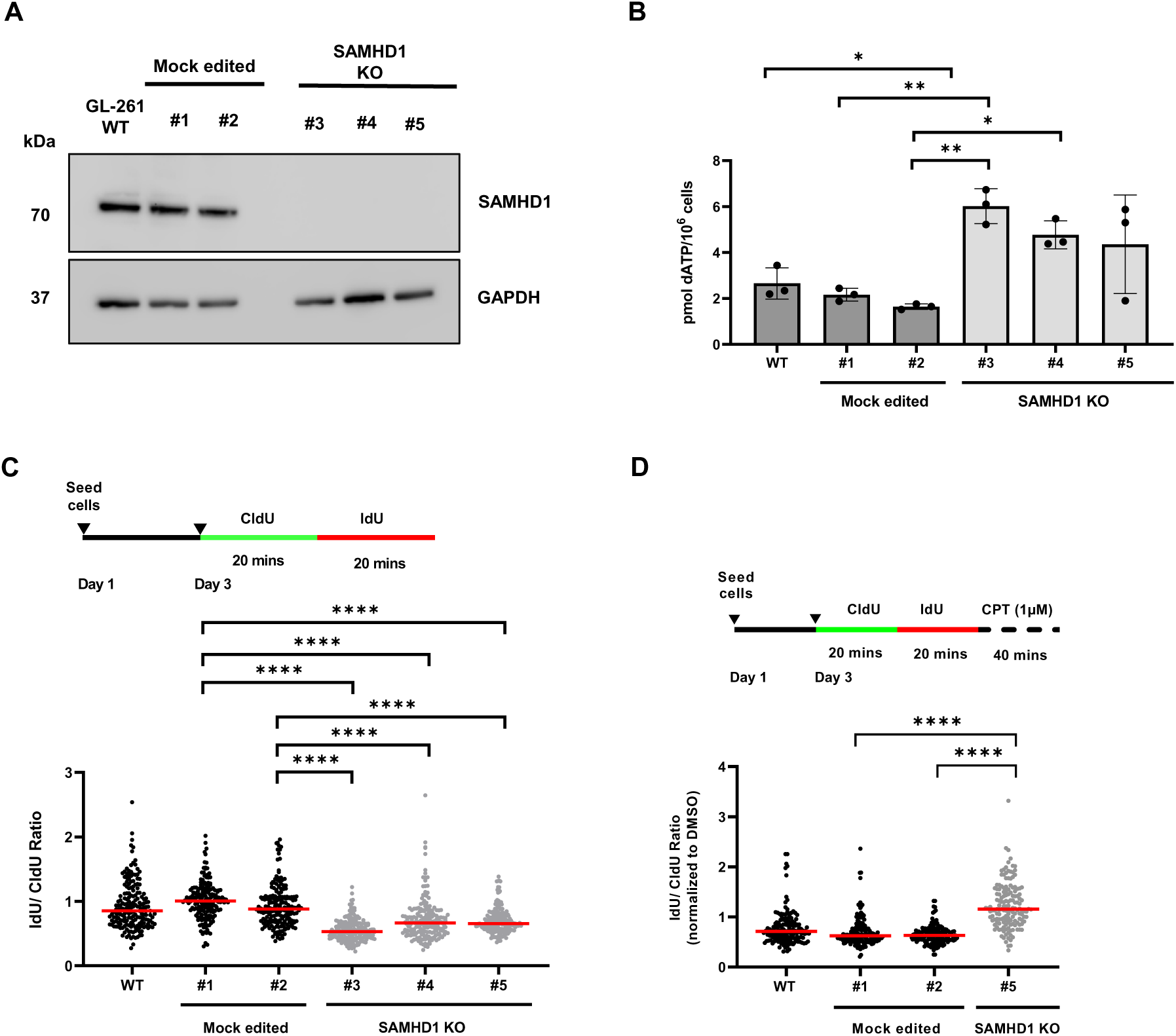
Murine SAMHD1 deficiency disrupts dNTP homeostasis, impairs replication fork dynamics and attenuates CPT-induced nascent DNA degradation. (A) SAMHD1 knock-out (KO) clones (samples #3, #4, and #5) and mock-edited clones (samples #1 and #2), were analyzed via immunoblot for SAMHD1 expression (mAB anti-SAMHD1 OTI3F5, Novus). GAPDH was used as a loading control. (B) Intracellular dATP levels were quantified in wild-type (WT), mock-edited and SAMHD1 KO GL261 cells using a single-nucleotide incorporation assay. Measurements were performed in three independent experiments. Bar graphs indicate mean of experiments, error bars correspond to standard deviation. Statistical significance was determined among all pairs of samples using Ordinary one-way ANOVA (Tukey’s multiple comparisons test) (*P < 0.05, **P < 0.01). (C) DNA fiber analysis was performed to assess unperturbed replication fork progression in proliferating WT, mock-edited, and SAMHD1 KO GL261 cells. Replication dynamics were evaluated by calculating the IdU/CldU tract-length ratio. Each dot represents an individual DNA fiber, and red horizontal lines indicate the median. A total of 188 replication forks were analyzed per sample (two slides for each condition). Statistical significance was determined using the Kruskal–Wallis test followed by Dunn’s multiple-comparisons test (****P < 0.0001). (D) Following sequential incubation with CldU and IdU, cells were treated with 1 µM camptothecin (CPT) for 40 min to induce replication-associated DNA damage. CPT-induced shortening of the IdU tract was assessed by calculating the IdU/CldU ratio, normalized to the corresponding DMSO-treated control. Each dot represents an individual DNA fiber, and red horizontal lines indicate the median. A total of 153 replication forks were analyzed per sample (two slides for each condition). Statistical significance was determined using the Kruskal–Wallis test followed by Dunn’s multiple-comparisons test (****P < 0.0001).

Since hSAMHD1 has been implicated in DNA replication and replication fork maintenance (20), we next investigated whether mSAMHD1 deficiency affects DNA replication dynamics in murine cells. Previous studies have associated SAMHD1 deficiency with altered dNTP homeostasis and impaired replication dynamics in human cells (26, 27). We therefore performed DNA fiber assays using sequential incorporation of the thymidine analogues CldU and IdU (28). Replication dynamics were assessed by determining the IdU/CldU tract-length ratio, with values approaching 1 indicating comparable DNA synthesis during the two labeling periods. SAMHD1-KO GL261 cells exhibited significantly reduced IdU/CldU ratios, whereas wild-type and mock-edited cells showed ratios close to 1 (Fig. 2C; Supplementary Fig. 2A). The shorter IdU tracts relative to CldU tracts indicate reduced DNA synthesis during the second labeling period relative to the first, consistent with impaired replication fork progression.

**Figure 2:**
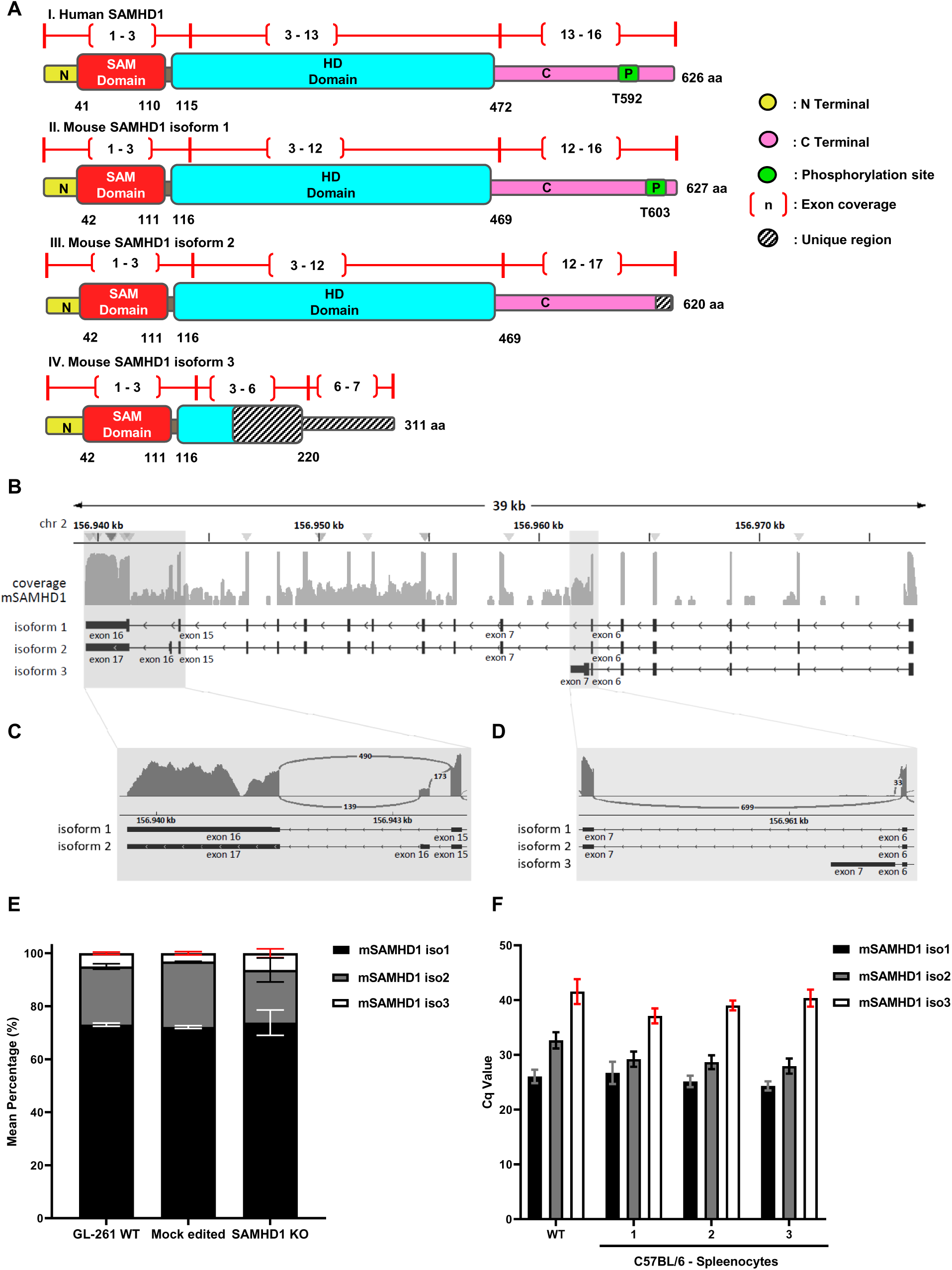
RNA sequencing reveals distinct murine *Samhd1* transcript isoforms in GL261 cells. (A) Schematic representation of human SAMHD1 (hSAMHD1) and the three annotated murine SAMHD1 (mSAMHD1) protein isoforms. Protein structures are shown according to NCBI RefSeq annotations. P indicates the corresponding C-terminal phosphorylation sites T592 in hSAMHD1 and T603 in mSAMHD1 isoform 1 (iso1). RefSeq accession numbers: hSAMHD1, NM_015474.4; mSAMHD1 iso1, NM_018851.4; iso2, NM_0011395202.2; and iso3, NM_001370610.1. (B) Alternative splicing of *mSamhd1* transcripts in GL261 cells. Representative RNA-sequencing coverage tracks and Sashimi plots illustrate exon coverage, splice junctions, and corresponding junction-spanning read counts. Plots were generated using Integrative Genomics Viewer (IGV). (C) Alternative splicing distinguishes *mSamhd1* iso2 from iso1. Isoform-specific splicing was assessed using junction-spanning reads connecting exons 15–16 (iso1) and exons 15–16– 17 (iso2). (D) A distinct alternative terminal exon distinguishes *mSamhd1* iso3 from iso1 and iso2. Iso3-specific splicing was identified by reads spanning the exon 6–7* junction, compared with the exon 6–7 junction shared by iso1 and iso2. (E) Relative abundance of *mSamhd1* isoform-associated splice-junction reads in GL261 cells. Mean percentages were determined in data sets of wild-type (WT, N=3), mock-edited (N=2), and SAMHD1-knockout (KO, N=5) GL261 cells. Values are presented as mean ± SD (F) RT-qPCR analysis of *mSamhd1* isoforms in GL261 cells (N=3) and primary C57BL/6 splenocytes (N=2) in three technical replicates. Mean quantification cycle (Cq) values were determined using isoform-specific RT-qPCR assays. Values are presented as mean ± SD.

hSAMHD1 has been implicated in DNA end resection and the processing of stalled replication forks (9, 10). In particular, SAMHD1 promotes MRE11-dependent degradation of nascent DNA at stalled replication forks (10). To investigate whether mSAMHD1 contributes to this process, cells were sequentially labeled with CldU and IdU. Replication stalling was induced using camptothecin (CPT), which inhibits topoisomerase I, to assess the stability of nascent DNA at stressed replication forks. DMSO-treated cells served as controls. Following CPT treatment, wild-type and mock-edited GL261 cells exhibited a significantly greater reduction in the IdU/CldU ratio than SAMHD1-KO cells (Fig. 2D, Supplementary Fig. 2B-2F). The attenuated response in SAMHD1-KO cells therefore indicates that mSAMHD1 contributes to CPT-induced nascent DNA processing suggesting a role in DNA end resection.

Together, these results demonstrate that SAMHD1 deficiency in murine GL261 cells is associated with elevated intracellular dATP levels, impaired replication fork progression, and reduced CPT-induced nascent DNA degradation, supporting a role for mSAMHD1 in nucleotide homeostasis and replication fork maintenance.

### RNA sequencing reveals differential expression of SAMHD1 isoforms in murine cells

Three murine *Samhd1* (*mSamhd1)* isoforms are currently annotated in the NCBI RefSeq database, and their predicted protein structures are illustrated in Fig. 2A. To characterize *mSamhd1* isoform expression in GL261 cells, we performed RNA sequencing and visualized alternative splicing using Sashimi plots generated with Integrative Genomics Viewer (IGV).

Analysis of splice-junction reads revealed distinct splicing patterns corresponding to the annotated *mSamhd1* isoforms (Fig. 2B). Isoform 2 (iso2) was distinguished from isoform 1 (iso1) by alternative splicing involving exons 15–17 (Fig. 2C; Supplementary Fig. 3A), whereas isoform 3 (iso3) was identified by its distinct terminal exon 7 (Fig. 2D; Supplementary Fig. 3B). Quantification revealed the highest expression of iso1, followed by iso2 and iso3 (Fig. 2E; Supplementary Fig. 3C). In wild-type GL261 cells, iso1 expression was approximately threefold higher than iso2 and 15-fold higher than iso3 (mean estimated read counts for *mSamhd1* isoforms in GL261 WT: iso1: 414.46 ± 100.83; iso2: 125.54 ± 37.47; iso3: 28.0 ± 4.58). Similar ratios were observed in mock-edited and SAMHD1-KO cells.

To examine whether this expression pattern also occurs in primary murine cells, we analyzed splenocytes from C57BL/6 mice by RT-qPCR (Fig. 2F, Supplementary Fig. 3D). Consistent with the RNA-sequencing results, iso1 exhibited the lowest Cq values, followed by iso2 and iso3. Together, these findings indicate that iso1 is the predominant *mSamhd1* transcript in murine cells.

### Murine SAMHD1 isoforms are enzymatically active and restore replication fork dynamics and nascent DNA processing in SAMHD1-deficient human cells

To discern the cellular functions of the different mSAMHD1 isoforms and compare with human SAMHD1 (hSAMHD1), we expressed FLAG-tagged mSAMHD1 isoforms in SAMHD1-knockout (KO) HEK293T cells. Immunoblot analysis revealed comparable expression levels of mSAMHD1 iso1 and iso2, whereas iso3 showed markedly lower expression (Fig. 3A, upper panel). Consistent with the absence of the phosphorylation site T603 in iso2 (Fig. 2A), iso1 and iso2 could be distinguished using a phospho-SAMHD1 antibody (Fig. 3A, lower panel). Immunofluorescence analysis further revealed predominantly nuclear localization of both iso1 and iso2 (Fig. 3B), consistent with the reported nuclear localization of hSAMHD1 (29).

**Figure 3:**
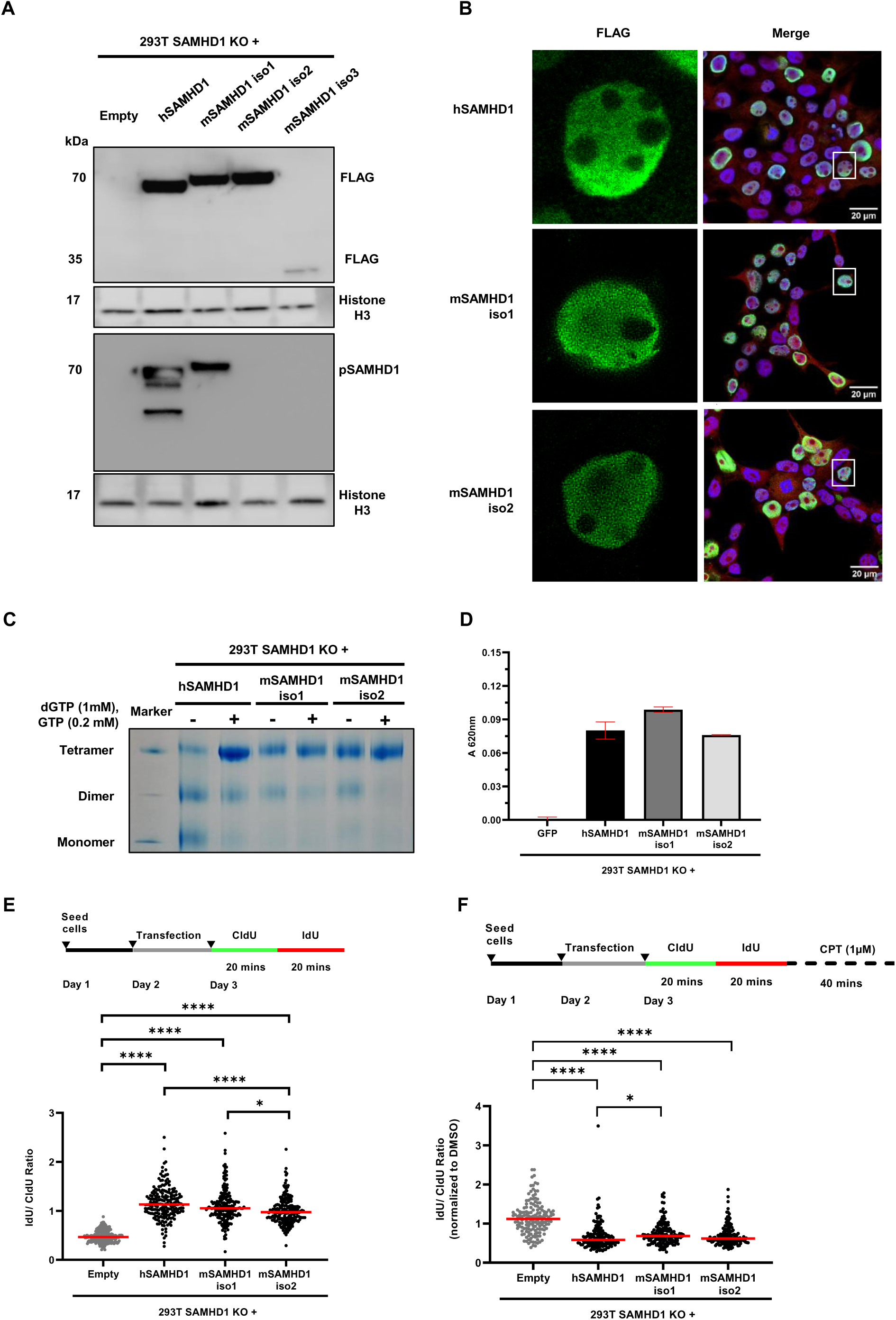
Murine SAMHD1 isoforms 1 and 2 restore replication-associated functions in SAMHD1-deficient human cells. (A) Representative immunoblot of FLAG-tagged human SAMHD1 (hSAMHD1) and murine SAMHD1 (mSAMHD1) isoforms expressed in HEK293T SAMHD1-knockout (KO) cells. Protein expression was detected using an anti-FLAG antibody, and phosphorylation was assessed using a phospho-SAMHD1 (pSAMHD1) antibody recognizing the human T592 phosphorylation site. Histone H3 served as a loading control. (B) Representative immunofluorescence images of HEK293T SAMHD1-KO cells transfected plasmids encoding hSAMHD1 or mSAMHD1 isoforms. FLAG-tagged SAMHD1 was detected by immunofluorescence, nuclei were counterstained with Hoechst, and the cytoplasm was visualized using CellMask. Individual fluorescence channels and merged images are shown. Images were acquired at 93× magnification using glycerol immersion. (C) FLAG-tagged hSAMHD1, mSAMHD1 isoform 1 (iso1), and mSAMHD1 isoform 2 (iso2) were affinity-purified from transfected HEK293T SAMHD1-KO cells using anti-FLAG M2 affinity gel to analyze nucleotide-dependent oligomerization of human and murine SAMHD1. Purified proteins were incubated with 1 mM dGTP and 0.2 mM GTP for 15 min at room temperature and subsequently analyzed by native gel electrophoresis to assess oligomerization (representative image). (D) Enzymatic activity of affinity-purified hSAMHD1, mSAMHD1 iso1, and mSAMHD1 iso2 was assessed using a malachite green-based phosphate detection assay. Samples were incubated with GTP and dGTP at 30 °C for 30 min. Phosphate release was quantified by measuring absorbance at 620 nm. Values are presented as mean ± SD of three technical replicates. (E) DNA fiber analysis was performed in SAMHD1-KO cells and SAMHD1-KO cells expressing hSAMHD1, mSAMHD1 iso1, or mSAMHD1 iso2. Replication dynamics were evaluated using the IdU/CldU tract-length ratio. Each dot represents an individual replication fork, and red horizontal lines indicate the median. A total of 200 replication forks were analyzed per sample (two slides for each condition). Statistical significance was determined using the Kruskal–Wallis test followed by Dunn’s multiple-comparisons test (*P < 0.05, ****P < 0.0001). (F) DNA fiber analysis was performed in HEK293T SAMHD1-KO cells, and SAMHD1-KO cells expressing hSAMHD1, mSAMHD1 iso1, or mSAMHD1 iso2. Following sequential CldU and IdU incubation, replication stress was induced by treatment with 1 µM camptothecin (CPT). CPT-induced nascent DNA degradation was assessed using IdU/CldU tract-length ratios normalized to the corresponding DMSO-treated controls. Each dot represents an individual replication fork, and red horizontal lines indicate the median. A total of 150 replication forks were analyzed per sample (two slides for each condition). Statistical significance was determined using the Kruskal–Wallis test followed by Dunn’s multiple-comparisons test (*P < 0.05, ****P < 0.0001).

Previous biochemical studies demonstrated nucleotide-dependent oligomerization of mSAMHD1 (20). To examine this property in our ectopic cellular expression system, FLAG-tagged proteins were affinity-purified and analyzed by native gel electrophoresis following incubation with GTP and dGTP. Both iso1 and iso2 exhibited a shift from predominantly dimeric species toward tetrameric complexes upon nucleotide addition (Fig. 3C). Furthermore, both purified isoforms from mammalian cells displayed dNTPase activity (Fig. 3D, Supplementary Fig. 4A) as measured with a malachite green colorimetric assay.

We next investigated whether mSAMHD1 isoforms could restore DNA replication dynamics in SAMHD1-deficient human cells. DNA fiber analysis revealed reduced IdU/CldU tract-length ratios in SAMHD1-KO HEK293T cells. Expression of hSAMHD1 or the investigated mSAMHD1 isoforms significantly increased these ratios to approximately 1, indicating restoration of replication fork progression (Fig. 3E, Supplementary Fig. 4B and 4C). While the rescue mediated by mSAMHD1 isoform 1 was comparable to that of hSAMHD1, a small but statistically significant difference was observed between the two murine isoforms. Nevertheless, both mSAMHD1 isoforms effectively restored the IdU/CldU tract-length ratios in the SAMHD1-null background.

Finally, we assessed CPT-induced nascent DNA degradation using DNA fiber analysis. Following CPT treatment, SAMHD1-KO HEK293T cells retained longer IdU tracts than cells with ectopic expression of hSAMHD1, consistent with reduced nascent DNA degradation. Expression of both mSAMHD1 isoforms restored CPT-induced IdU tract shortening to levels comparable to those observed following hSAMHD1 expression (Fig. 3F; Supplementary Fig. 4D, 4E, 4F).

Together, these results demonstrate that mSAMHD1 iso1 and iso2 share key biochemical and cellular properties with hSAMHD1, including nuclear localization, nucleotide-dependent oligomerization, dNTPase activity, and the ability to restore replication fork dynamics and CPT-induced nascent DNA processing in SAMHD1-deficient human cells.

## Discussion

Human SAMHD1 (hSAMHD1) regulates intracellular dNTP homeostasis, restricts HIV-1 replication, contributes to DNA replication and repair, and modulates innate immune signaling (2). Although murine SAMHD1 (mSAMHD1) has been characterized with respect to its dNTPase activity, antiviral restriction, and regulation of innate immune responses (17, 16, 22, 24, 21), its contribution to DNA replication dynamics and DNA end resection remains less well understood. In this study, we investigated these functions in SAMHD1-deficient murine GL261 glioma cells and compared the cellular activities of individual mSAMHD1 isoforms with hSAMHD1.

SAMHD1 knockout in GL261 cells resulted in elevated intracellular dATP levels, impaired replication fork progression, and reduced CPT-induced nascent DNA degradation. These findings are consistent with previous observations in human cells, where SAMHD1 deficiency has been associated with altered dNTP homeostasis and replication dynamics (26, 27). Furthermore, Coquel et al. (2018) demonstrated that hSAMHD1 promotes MRE11-dependent processing of stalled replication forks and contributes to replication fork restart (10). Our findings extend these observations to murine cells and support a conserved role of SAMHD1 in replication fork maintenance. However, the relative contributions of altered nucleotide homeostasis and other SAMHD1-dependent mechanisms to the observed replication defects remain to be determined. Likewise, although the reduced CPT-induced shortening of nascent DNA tracts is consistent with impaired fork-associated DNA degradation, further studies are required to establish whether mSAMHD1 directly regulates MRE11 activity or DNA end resection at double-strand breaks.

In addition to investigating mSAMHD1 function, we characterized the expression of its annotated splice isoforms. Previous studies described two major mSAMHD1 isoforms with differences in their C-terminal regions (21, 3). A third transcript variant is currently annotated in the NCBI RefSeq database (NM_001370610.1; NP_001357539.1), but its expression and cellular functions remain poorly characterized. Our RNA-sequencing analysis identified splice patterns corresponding to all three annotated isoforms in GL261 cells, with isoform 1 showing the highest estimated expression, followed by isoforms 2 and 3. This distribution is consistent with the previously reported predominance of isoform 1 in murine tissues (21). RT-qPCR analysis of murine splenocytes further supported the predominance of isoform 1. These findings suggest that all three annotated mSAMHD1 transcript variants are expressed in murine cells, although their relative abundance may vary between cell types.

We next investigated whether the individual mSAMHD1 isoforms share the cellular functions established for hSAMHD1. Previous studies demonstrated dNTPase activity and antiviral restriction by the two major murine isoforms, as well as nucleotide-dependent oligomerization (21, 20, 22). Consistent with these findings, mSAMHD1 isoforms 1 and 2 retained dNTPase activity and formed higher-order oligomeric complexes upon nucleotide addition in our mammalian cellular system. Both isoforms also localized predominantly to the nucleus, similar to hSAMHD1(29). Importantly, expression of either murine isoform in SAMHD1-deficient HEK293T cells restored replication fork dynamics and CPT-induced nascent DNA degradation to levels comparable to those observed following hSAMHD1 expression. These results provide evidence that the two major mSAMHD1 isoforms retain cellular functions associated with replication fork maintenance, despite differences in their C-terminal regions.

In contrast to isoforms 1 and 2, mSAMHD1 isoform 3 showed markedly reduced protein expression following transfection into HEK293T cells, reflecting the relative expression level in primary murine cells, limiting its functional characterization. Its truncated protein sequence and distinct C-terminal region may influence protein stability or expression. However, the underlying mechanism remains unknown, and the current findings do not distinguish between reduced transcript abundance, altered translation, and increased protein turnover. Further studies examining transcript stability, protein degradation, and expression under alternative experimental conditions will be required to determine whether isoform 3 produces a stable, functional protein.

Although isoforms 1 and 2 differ in their C-terminal regions, including the absence of the residue corresponding to the regulatory hSAMHD1 phosphorylation site T592 in isoform 2, both restored replication-associated functions in our experimental system. This suggests that the C-terminal differences between these isoforms do not prevent their activity in the assays investigated here. However, the present findings do not exclude isoform-specific differences in other cellular functions or under different regulatory conditions. In particular, whether phosphorylation-dependent regulation contributes to their activity during replication stress remains to be investigated.

Beyond its established roles in nucleotide metabolism and genome maintenance, SAMHD1 has been implicated in several human malignancies. Somatic SAMHD1 mutations have been identified in chronic lymphocytic leukemia and colorectal cancer, whereas altered SAMHD1 expression has been reported in lung adenocarcinoma. Beyond its potential involvement in tumor biology, SAMHD1 also influences responses to anticancer therapies, as demonstrated by its ability to confer resistance to the nucleoside analogue cytarabine in hematological malignancies. (30, 31, 13, 32). These observations suggest that SAMHD1 dysfunction may influence tumor biology, although its effects are likely to depend on tumor type and cellular context.

In glioblastoma, SAMHD1 expression has been associated with clinical outcomes and treatment responses. Liu et al. (2023) reported that elevated SAMHD1 expression was associated with poorer survival in chemotherapy-treated patients with glioblastoma and glioma (33). Furthermore, Daddacha et al. (2022) demonstrated improved survival following implantation of SAMHD1-deficient human LN-229 glioblastoma cells in immunodeficient mice (34). More recently, Sun et al. (2025) reported improved survival in an orthotopic glioma model following suppression of SAMHD1 and described changes in innate immune signaling in vitro (14). Together, these studies suggest that SAMHD1 may influence glioma progression and treatment responses. However, xenograft models using immunodeficient mice cannot fully capture interactions between tumor cells and an intact adaptive immune system. This limitation is particularly relevant for studying SAMHD1, given its role in regulating cytosolic nucleic acid sensing and inflammatory signaling, which may influence interactions between tumor cells and the immune microenvironment.

The SAMHD1-deficient murine GL261 cells established in this study therefore provide a potential experimental model for investigating SAMHD1 function in an immunocompetent, syngeneic glioma setting. Such a model could help determine how tumor-cell SAMHD1 influences tumor growth, responses to DNA-damaging therapies, and interactions with the immune microenvironment. Given the involvement of SAMHD1 in nucleotide homeostasis, replication stress, and innate immune regulation, these processes may be particularly relevant to its role in glioma biology.

In conclusion, our findings support a conserved role of human and murine SAMHD1 in replication fork dynamics and CPT-induced nascent DNA processing. Furthermore, we identify transcripts corresponding to three annotated murine SAMHD1 isoforms and demonstrate that isoforms 1 and 2 can restore replication-associated functions in SAMHD1-deficient human cells. The established GL261 SAMHD1-KO model provides a basis for further mechanistic studies and future investigations into the contribution of SAMHD1 to glioma progression and the tumor immune microenvironment.

## Materials and Methods

### Cell lines and cell culture

For CRISPR/Cas9-mediated SAMHD1 knock-out in HEK293T (ATCC No.: CRL-11268), cells were transfected with pSpCas9(BB)-2A-GFP (Addgene), where the gRNA targeting human SAMHD1 at exon 1 (5′-ATCGCAACGGGGACGCTTGG-3′) was cloned into the scaffold. After single cell cloning, clone #55 was verified by western blotting. To confirm bi-allelic SAMHD1 KO, the modified region was amplified using primers III-EX1_FW (5′-GAGCCTGCTCCCATCCTACGAA) and III-EX1_RV (5′-TTCTAGCCACGCCCTTTCGC) with GO Taq Polymerase (Promega), followed by cloning into pGEM T-easy vector system (Promega) and Sanger sequencing. The cells were cultured in DMEM (Sigma-Aldrich) supplemented with 10% fetal calf serum (FCS, Sigma-Aldrich) and 2mM L-glutamine (Sigma-Aldrich) at 37°C under 5% CO_2_. Murine GL261 cells (a kind gift of Mirko Schmidt, DSMZ no. ACC 80*2*) were cultured at 37°C under 8% CO_2_ in DMEM supplemented with 10% FCS and 2mM L-glutamine. All cell lines were free of mycoplasma contamination, as tested by PCR Mycoplasma Test Kit II (PanReac AppliChem).

### CRISPR/Cas9 knock-out of GL261 cells

200µM Edit-R Modified Synthetic crRNA targeting SAMHD1 exon 4 (Mm.Cas9.SAMHD1.1.AF with PAM site AGG, target sequence: 5′-TCGATGATTCTGATAAGGAG, Dharmacon), 200 µM Edit-R CRISPR-Cas9 Synthetic tracrRNA (Dharmacon) and 40µM Cas9-Nuclease Protein (QB3 Macrolab) were assembled *in vitro*, as previously described and optimized (35, 36). These ribonucleoproteins were added into 5 × 10^5^ sub-confluent GL261 cells using 4D-Nucleofector X Unit and SF Cell line Kit (Lonza) in DN-100 program. Single cell clones were generated using limited dilution after nucleofection. To confirm SAMHD1 KO, the modified region was amplified using primers mSAMHD1_E4_TIDE_fwd (10µM) (5′-GGT GTA AAA CAC AGG CAA ACA C) and mSAMHD1_E4_TIDE_rev (10µM) (5′-CAG TCA TAC CAG CAA CCA GC) together with GoTaq® DNA Polymerase (Promega), followed by TA cloning into pGEM T-easy vector system (Promega) and Sanger sequencing.

### dNTP quantification

1×10^6^ GL261 cells were washed with cold PBS and lysed with 200µl 65% cold methanol under vigorous vortex for 2 min. The lysate samples were centrifuged in a speed vacuum at 45°C until the supernatant methanol was evaporated. The quantification of dATP was performed using single nucleotide incorporation assay (25).

### DNA fiber spreading assay

The DNA fiber spreading assay was performed as previously described (28). Subconfluent HEK293T/GL261 cells were incubated with 20µM 5-chloro-2’-deoxyuridine (CldU, Sigma-Aldrich) and then with 200µM 5-iodo-2’-deoxyuridine (IdU, Sigma-Aldrich) at 37°C, 5% CO_2_ for exactly 20 min. 2500 cells were spotted on top of Superfrost microscopic slides (Erepdia) and gently resuspended in Lysis Buffer (0.5% (v/v) SDS, 200mM Tris-HCl (pH 7.4), 50mM EDTA in H_2_O) and allowed to lyse at room temperature (RT) for 6 min. The slides were tilted around 30° angle and allowed run along the glass slides, air dried for 6 min, fixed in 3:1 Methanol: Acetic acid at RT for 5 min, and allowed to air dry for 7 min. The samples were denatured in 2.5N HCl at RT for 1 hour. The glass slides were subsequently block in 5% BSA and immunostained with CldU cross reactive rat anti-BrdU and IdU cross reactive mouse anti-BrdU primary antibodies. Anti-rat AF488 and anti-mouse AF555 were correspondingly used. The fiber samples were mounted in Cover Glass (24×50 mm) using Mowiol-4-88 (Calbiochem)-DABCO (Roth) mix (3:1 ratio). For each sample, 150–250 DNA fibers containing both red and green tracks were imaged using a Zeiss Axio Observer 3 microscope equipped with a 63×/1.4 NA oil-immersion objective, and the lengths of both tracks were measured using ImageJ. Fiber assay was conducted on two independent slides and samples were randomized during analysis. Statistical analysis was performed with GraphPad Prism software using Kruskal-Wallis tests with Dunn’s nonparametric multiple comparisons tests.

### RNA sequencing and isoform quantification

We sequenced 6 different clones across 3 conditions: 1 WT clone (N=3, all replicates), 2 mock-edited clones (N=2 singletons) and 3 SAMHD1 KO clones (N=5, 3 replicates of one clone and 2 singleton clones), resulting in 10 RNAseq data sets in total. Capture of mRNA from 1.5×10^6^ GL261 cell pellets using the RNeasy Mini Kit (Qiagen), sequencing library preparation with the TrueSeq Stranded mRNA Library Prep Kit (Illumina, San Diego, USA) and sequencing on the NovaSeq6000 (S1 flow cell kit; Illumina, San Diego, USA) with paired end 2 x 150bp setting were performed by TRON gGmbH (Mainz, Germany). The quality control of RNA sequencing raw fastqs was performed using fastp v.0.20.0 (37). Default parameters were used with the following exceptions; a sequence for adapter trimming was specified, trimming from front (5’) to tail (3’) and vice versa based on mean quality scores in a sliding window was applied and polyG trimming was disabled (–– adapter_sequence =TCAGATCGGAAGAGCACACGTCTGA-ACTCCAGTCAC –– cut_front ––cut_tail-G). These quality controlled fastqs were then aligned with mouse reference genome GRCm39 (NCBI RefSeq assembly GCF_000001635.27R) using STAR v.2.7.11b. (38). The corresponding genomic annotation (.gtf) file was optimized for isoforms quantification (––chimSegmentMin 2 ––outFilterMismatchNmax 3 ––twopassMode Basic ––alignEndsType EndToEnd ––outSAMstrandField intronMotif –– alignSJDBoverhangMin 1 ––alignIntronMax 299999). The coverage and sashimi plots of the mSAMHD1 gene and its isoforms 1-3 (corresponding to RefSeq IDs NM_018851.4, NM_001139520.2 and NM_001370610.1) along with their corresponding splice junction-spanning read counts were generated using the “sashimi plot” function in IGV v.2.19.2 (39). RNAseq-based isoform abundances were extrapolated based on the splice junction-spanning read counts between the isoform-defining exons (Fig. 2C-E, Supplementary Fig. 3A-B). Mere read counts spanning exons 6-7 were used for isoform 3 abundances, but since isoform 1 and 2 are identical at this location, and coverage varies across the length of the sequence, their respective abundances were estimated by adjusting the combined read count across exons 6-7 based on their relative frequencies at exon 15-16 (isoform 1) and exon 15-17 (arithmetric mean of read counts across both junctions, isoform 2).

### qPCR-based quantification of mSAMHD1 isoforms

Total RNA was extracted from GL261 or C57BL/6 mice splenocytes using Direct-zol-96 RNA (Zymo Research), followed by quantitative real-time PCR using QuantiTect® SYBR® Green RT-PCR kit (Qiagen) with primers for mSAMHD1 isoform specific region on CFX384TM Real-Time System (Bio-Rad). mSAMHD1 iso1 (21): (FW: 5’-CAAGCCACAGGATGGTGACA, RV: 5’-TCTTGGAGGCAAGATGAAGTCT), mSAMHD1 iso2 (21): (FW: 5’-CAAGCCACAGCAATGTGGGGCT, RV: 5’-CTGGTTGTGAGCCGCATGT), mSAMHD1 iso3: (FW: 5’-AAGTGGAAGCATCCTTCCCTG, RV: 5’-CTCCACAGGCTACTGTGAGGG) and GAPDH (16): (FW: 5’-AAGGGGCGGAGATGATGAC, RV: 5’-GGTGCTGAGTATGTCGTGGAG), mGAPDH: (FW: 5’-AAG GGG CGG AGA TGA TGA C; RW: 5’-GGT GCT GAG TAT GTC GTG GAG).

C57BL/6 mice were bred and housed in the animal facility of Paul-Ehrlich-Institut in accordance to standard procedures. All animal experiments were performed in accordance with the regulations of the German animal protection law and the respective European Union guidelines.

### Cloning of mSAMHD1 isoform constructs

mSAMHD1 isoform 1 construct has been described earlier (40). Open reading frame (ORF) clone for mSAMHD1 isoform 2 (OMu84857) and isoform 3 (OMu84855) were obtained from GenScript, USA as lyophilized plasmid DNA. By designing forward primers (10µM) with BamHI restriction site and FLAG tag and reverse primer (10µM) with EcoRI restriction site, PCR was conducted on the plasmids using KAPA HiFi HotStart ReadyMix (2X) (Roche) in T-Professional PCR-Thermocycler (Biometra) and ligated into pc DNA 3.1 (+) empty vector. mSAMHD1 iso 2: FW: 5’-GCGC GGA TCC ATG GAC TAC AAA GAC GAT GAC GAC AAG GGG CAG AGC GCA CCC TTG GAG, RV: 5’-GCGC GAA TTC TTA CAG CTG GTT GTG AGC CGC and mSAMHD1 iso3: FW:5’-GCGC GGA TCC ATG GAC TAC AAA GAC GAT GAC GAC AAG GGG CAG AGC GCA CCC TTG GAG, RV:5’-GCGC GAA TTC TTA AAA GAA AAC AAA GTC TAC AGT AAT CTG ACA GAG TAG AAC GGA.

### Intracellular localization

HEK293T SAMDH1 KO cells were transfected with 2µg of hSAMHD1/mSAMHD1 isoform plasmids using Lipofectamine 2000 (Invitrogen) (DNA vector: Lipofectamine ratio 1:4) for 48 hours. Cells were fixed using 3.7% PFA-PBS at RT for 10 min and permeabilized using 0.5% TritonX100-PBS at RT for 20 min. After blocking with 2% BSA-PBS for 30 min, the cover slips were immunostained with anti-FLAG mouse antibody (1:500, Sigma), Hoechst (1:10000, Invitrogen) and CellMask Deep Red (1:30000, Invitrogen). The corresponding secondary antibody AF 488 Goat anti-mouse (1:500, Invitrogen) was used for anti-FLAG.

### Immunoblot

Cells were collected, washed with PBS and lysed in radioimmunoprecipitation buffer [RIPA; 2mM EDTA, 1% glycerol, 137mM NaCl, 1% NP40, 0.1% SDS, 0.5% sodium deoxycholate, 25mM Tris (pH 8.0)] supplemented cOmplete Protease Inhibitor Cocktail [Roche] and PhosSTOP Phosphatase Inhibitor Cocktail [Roche]) on ice for 30 min. After centrifugation, the lysate solution was collected and measured for protein concentration using Protein Assay Dye Reagent Concentrate (BioRad) at 595 nm in a UV-Vis spectrophotometer (Harvard Bioscience). Using NuPAGE LDS Sample Buffer and Reducing Reagent (Thermo Fisher), 20 µg of protein was denaturated at 70°C for 10 min. The samples were ran on pre-cast 4-12% NuPAGE Bis-Tris gradient gel (Thermofisher) with PS10 plus protein marker (GeneOn BioScience) under MOPS running buffer (1M MOPS, 1M Tris, 69.3mM SDS, 20.5mM EDTA Titriplex II). After separation, the gels were placed on X Cell II Blot Module (Thermofisher) with methanol activated Hybond P 0.45 PVDF membrane (GE Healthcare). The PVDF membranes were blocked in 5% BSA or milk powder (Carl Roth) in TBST (Tris-buffered saline, 0.1% Tween; RT, 40 min). The incubation of primary antibodies was performed at 4°C, overnight, anti-GAPDH (14C10, CST), Histone H3 (MABI-0301) (39763, Active Motif, 1:1000), anti-FLAG M2 (F1804-200UG,1:2000), anti-SAMHD1 (12586-1-AP, Proteintech, 1:2000), anti-SAMHD1 OTI3F5 (NBP2-03285, Novus, 1:2000), and anti-SAMHD1-pT592 (D702M, CST, 1:1000) diluted in 5% BSA or milk powder in TBST. After washing in TBST, the membranes were incubated in horseradish peroxidase (HRP)-linked anti-rabbit IgG or anti-mouse IgG antibody (CST) at RT for 1 hour. The membranes were finally washed again before detection on a FUSION FX7 (Vilber Lourmat) using ECL Prime reagent (GE).

### FLAG tag pulldown and native gel electrophoresis

HEK293T SAMHD1 KO cells were transfected with hSAMHD1/mSAMHD1 isoform plasmids using Polyethylenimine (PEI, Sigma-Aldrich) (DNA vector: PEI ratio 1:3) for 48 hours. The cells were washed with cold PBS and lysed using NET lysis buffer (50mM Tris-HCl [pH 7.4], 150mM NaCl, 15mM EDTA [pH 7.4], 1% (v/v) NP40) at 4°C for 30 min. The lysates were incubated with 500ng/µl of ANTI-FLAG® M2 Affinity Gel beads (Sigma) under constant rotation at 40 rpm, 4°C for 1 hour. The beads were washed with NENT 300 buffer (20mM Tris-HCl [pH 7.4], 300mM NaCl, 1mM EDTA [pH 7.4], 0.1% (v/v) NP40, 25% (v/v) glycerol) and eluted using 500ng/µl N FLAG peptide (MedChemExpress, 63906) in TBS under constant rotation at 40 rpm, 4°C for 30 min. The eluted samples were incubated 1mM of dGTP (ThermoScientific) and 0.2mM of GTP (Thermo-Scientific) at RT for 15 min.

NativePAGE™ Sample Buffer (4X) (ThermoScientific) was added to the eluted samples. The samples were added to NativePAGE ™ Gel (ThermoScientific) in electrophoresis chamber (ThermoScientific). The electrophoresis was conducted at 150V for 60 min first and increased to 250V for another hour. After electrophoresis the gel was stained using 1X Problue Safe Coomassie Stain (Giotto Biotech) and imaged.

### Malachite green based dNTPase activity

As mentioned above, hSAMHD1 or mSAMHD1 was eluted with 500ng/µl N FLAG peptide in HEPES buffer (20mM HEPES pH 7.5, 150mM NaCl, 2.5mM MgCl_2_, 1mM DTT, 10% Glycerol). 0.1mg/ml hSAMDH1 or mSAMHD1 protein was incubated with 0.1mM dNTP mix at 30°C for 30 min. Malachite Green working reagent was added and incubated at RT for 30 min, as described in Malachite Green Phosphate Assay Kit (MAK307, Sigma-Aldrich). Using a microplate reader (TECAN-Spark^®^, Tecan), the absorbance (y) was measured at 620nm using TECAN software.

## Statistical Analysis

All statistical analysis were performed using GraphPad Prism (V9).

## Acknowledgements

We would like to acknowledge Francis Engelhardt, Ulrike Schumann and Mirko Schmidt for collaborative discussion and providing GL216 cells; Florian Krach for helpful discussions and Feyza Durak, Lisa Vetterdietz and Saskia Mönch for technical assistance. The project was funded by the Deutsche Forschungsgemeinschaft (DFG, German Research Foundation) – Project-ID 318346496 – SFB 1292 TP04 to R.K., Project-ID 450627322 – SFB 1506 B03 to M.R.-S. and L.W and National Institute of Health (NIH) R01 AI136581 and R01 AI162633 to B.K. Author contributions: conceptualization, B.M., D.R., N.V.F, R.K.; formal analysis, B.M., D.R., A.J., N.V.F., M.H., R.K., N.V.F.; investigation, B.M., D.R., A.J., N.V.F., M.H., K.S., F.D.H., Q.L., C.M., S.M., L.V., A.H.; resources, L.W., M. R.S., B.K., R.K.; writing—original draft preparation, B.M., N.V. F., D.R., A.J., R.K.; writing—review and editing, B.M., M.H., D.R., A.J., N.V.F., R.K.; visualization, B.M., D.R., A.J., M.H., N.V.F.; supervision, D.R., A.J., N.V.F., R.K.; funding acquisition, R.K., M.R.-S., L.W., B.K.

## Figure legends

**Supplementary figure 1:**
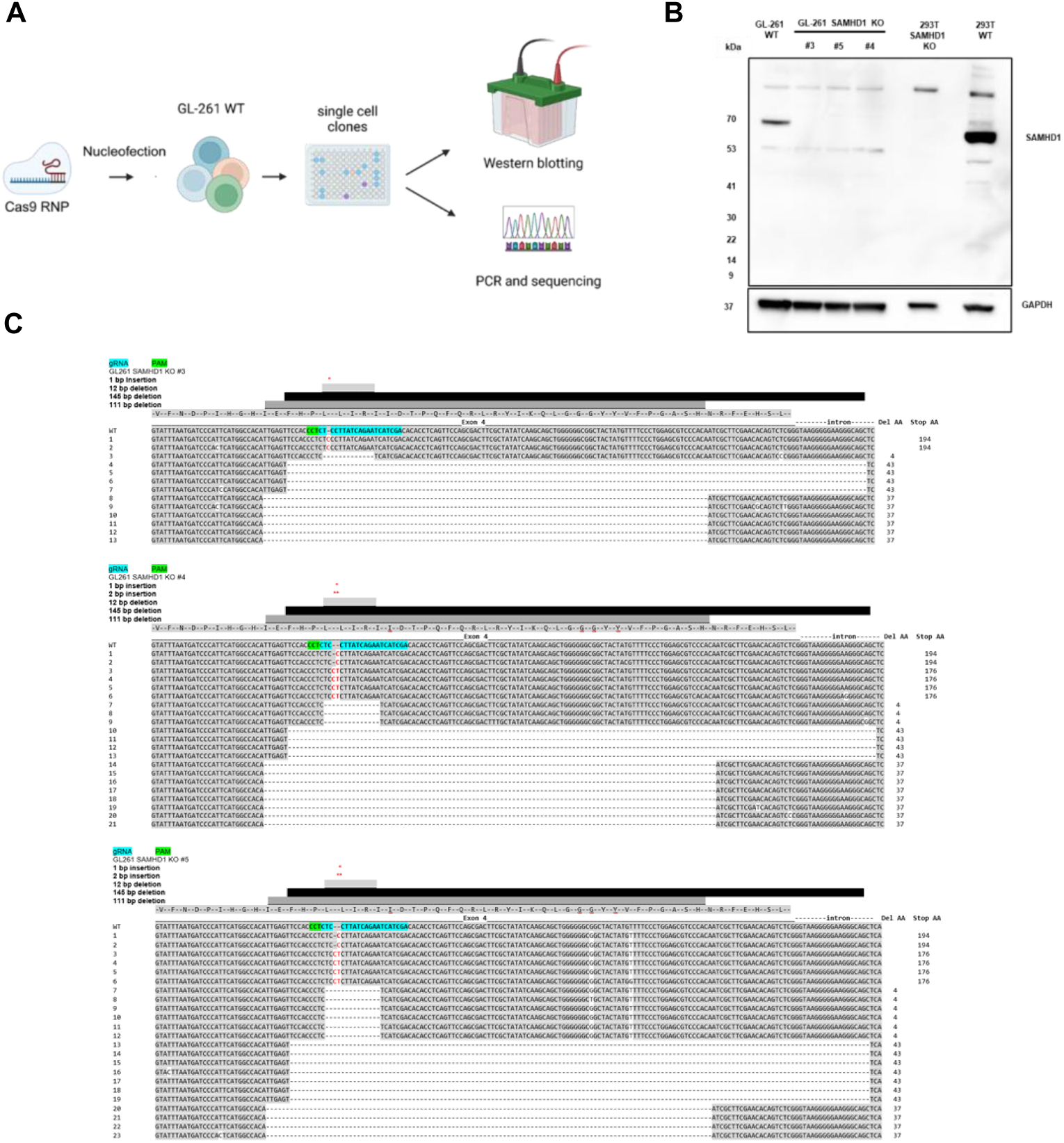
Generation of SAMHD1 knockout (KO) GL261 cells. (A) Schematic workflow of experimental steps and evaluation of CRISPR/Cas9 meditated knock-out by Western Blot and PCR based TA cloning and sequencing. Created with BioRender.com. (B) SAMHD1 knock-out (KO) GL261 clones (samples #3, #4, and #5), WT GL261 cells, 293T and SAMHD1 KO 293T cells as controls for protein loss, were analyzed via immunoblot for SAMHD1 expression (pAb, Proteintech, 12586-1-AP). GAPDH was used as a loading control. (C) Single cell clones were Sanger sequenced after TA-cloning to separate alleles and aligned to WT. Results of sequencing of SAMHD1 KO #3, #4 and #5 are depicted. Insertions (*, Red) and deletions (Grey or Black) of nucleotide base pairs are represented. gRNA sequence and PAM site position are indicated. Amount of amino acids deleted (AA del) and position of premature stop codon (Stop AA) are indicated.

**Supplementary Figure 2:**
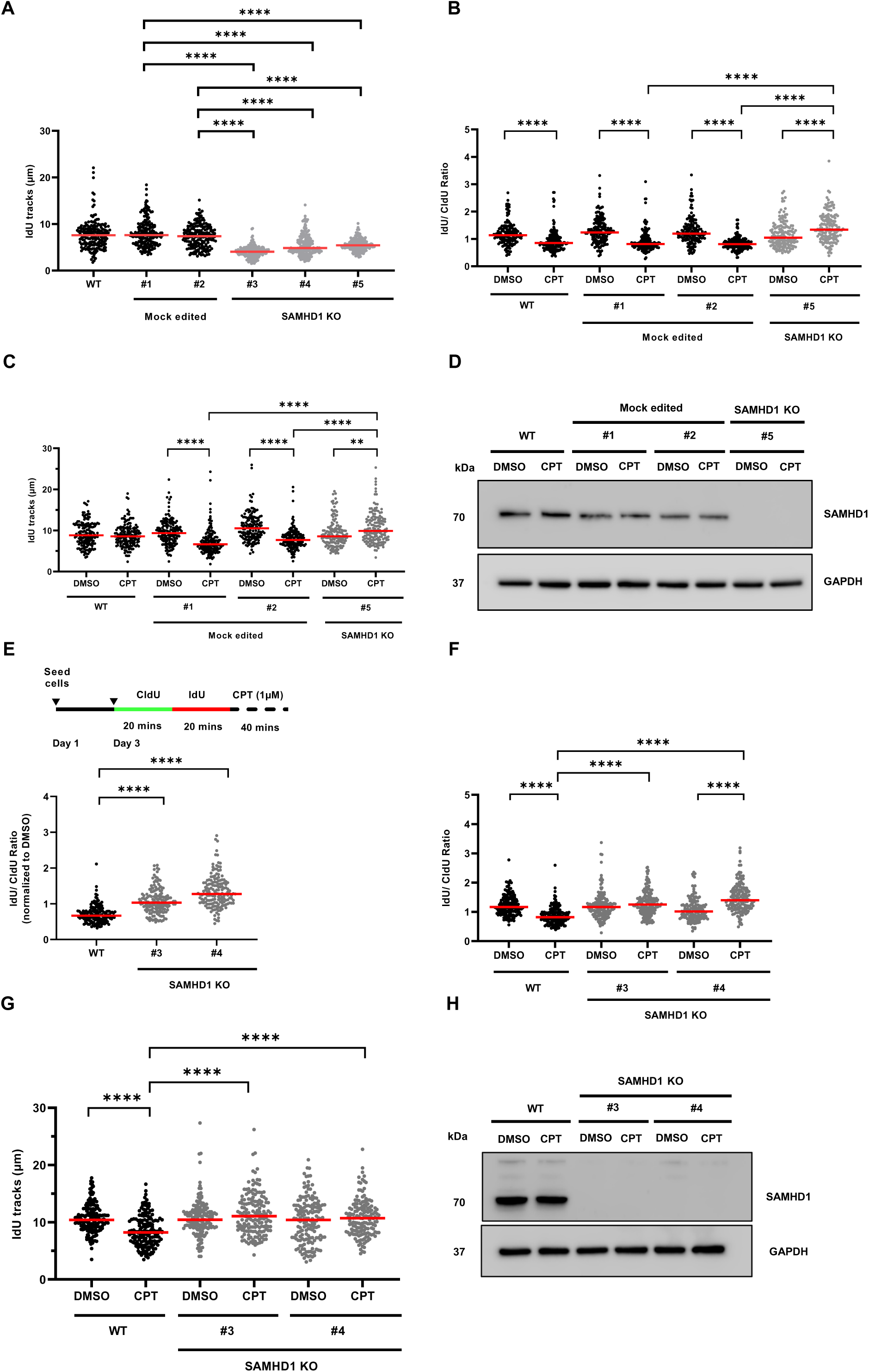
Impaired CPT-induced nascent DNA degradation in SAMHD1-knockout GL261 cells. (A) IdU track-lengths of DNA fiber analysis (188 replication forks, two slides for each condition) corresponding to Fig. 1C. Statistical significance was determined using the nonparametric Kruskal–Wallis test followed by Dunn’s multiple-comparisons test (****P < 0.0001). IdU/CldU tract-length ratios without normalization (B) and IdU track-lengths (C) of DNA fiber analysis (153 replication forks, two slides for each condition) corresponding to Fig. 1D. Statistical significance was determined using the nonparametric Kruskal–Wallis test followed by Dunn’s multiple-comparisons test (**P < 0.01, ***P < 0.0001). (D) Representative immunoblot corresponding to Fig. 1D and Supplementary Fig. 2B and 2C. SAMHD1 was detected using an anti-SAMHD1 antibody (OTI3F5, mouse monoclonal, Novus Biologicals). GAPDH served as a loading control. DNA fiber analysis of CPT-induced nascent DNA degradation or DMSO-control treatment, expressed as IdU/CldU track-length ratios and IdU track-lengths. CPT-induced nascent DNA degradation in additional SAMHD1-knockout GL261 samples. IdU/CldU tract-length ratios are shown (E) normalized to the corresponding DMSO-treated controls and (F) without normalization with (G) individual IdU track-lengths. A total of 161 replication forks were analyzed per sample (two slides for each condition). Statistical significance was determined using the nonparametric Kruskal–Wallis test followed by Dunn’s multiple-comparisons test (****P < 0.0001). (H) Immunoblot corresponding to Supplementary Fig. 2E, 2F and 2G. SAMHD1 was detected using an anti-SAMHD1 antibody (OTI3F5, mouse monoclonal, Novus Biologicals. GAPDH served as a loading control.

**Supplementary Figure 3.**
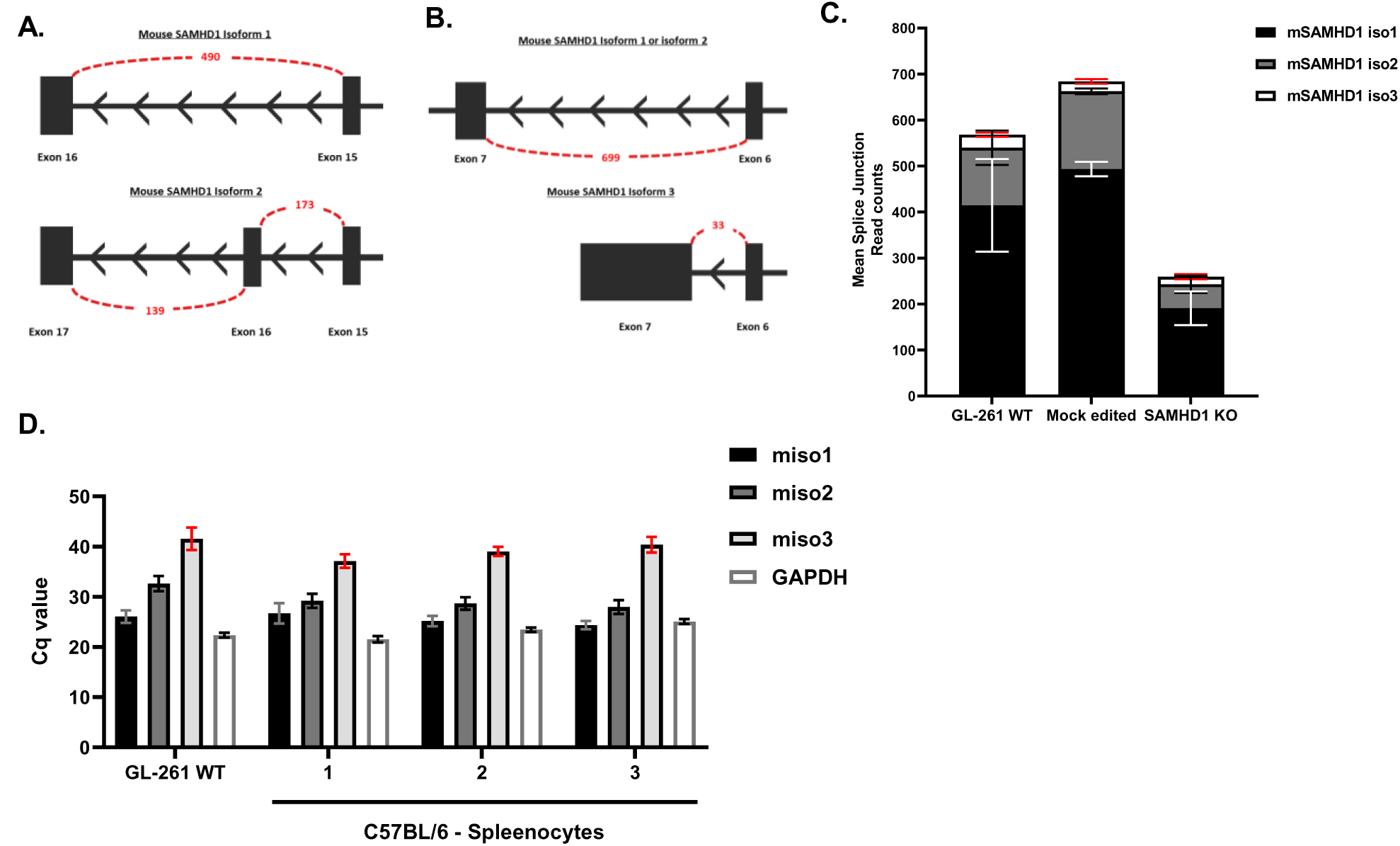
Splice-junction read counts and quantification of murine *Samhd1* isoforms in GL261 cells. (A) Distinction between *mSamhd1* isoforms 1 and 2 in GL261 wild-type (WT) cells. Isoform-specific splice junctions were identified based on the additional exon 17 in *mSamhd1* isoform 2. Junction-spanning reads connecting exons 15–16 and 16–17 were used to distinguish the two isoforms, as illustrated schematically. (B) Distinction between *mSamhd1* isoforms 1/2 and isoform 3 in GL261 WT cells. Isoform 3 contains a distinct terminal exon 7, allowing its identification through isoform-specific splice-junction reads. Since isoforms 1 and 2 share the same exon structure in this region, they cannot be distinguished using these junctions alone. (C) Splice-junction read counts associated with *mSamhd1* isoforms in GL261 cells. Mean junction-spanning read counts were estimated for *mSamhd1* isoforms 1, 2, and 3 in WT, mock-edited, and SAMHD1-knockout (KO) GL261 cells. (D) RT-qPCR analysis of *mSamhd1* isoforms related to Figure 2F. Mean quantification cycle (Cq) values were determined using isoform-specific RT-qPCR assays including house-keeping gene GAPDH. Values are presented as mean ± SD.

**Supplementary Figure 4.**
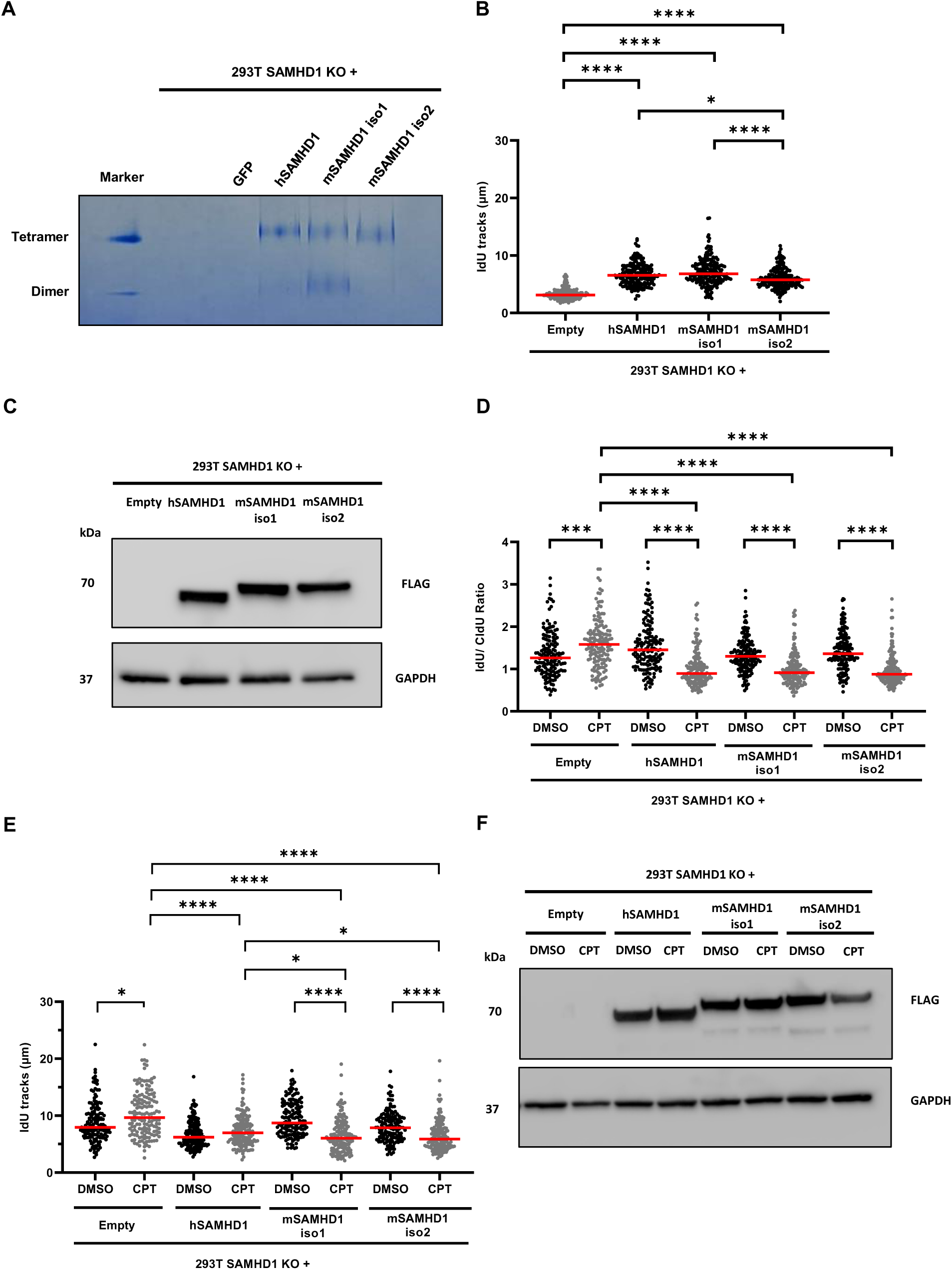
Murine SAMHD1 isoforms 1 and 2 restore CPT-induced nascent DNA degradation in SAMHD1-deficient HEK293T cells. (A) Representative native gel electrophoresis cooresponding to Fig. 3D. (B) IdU track-lengths of DNA fiber analysis (200 replication forks, two slides for each condition) corresponding to Fig. 3E. Statistical significance was determined using the nonparametric Kruskal–Wallis test followed by Dunn’s multiple-comparisons test (*P < 0.05, ****P < 0.0001). (C) Representative immunoblot corresponding to Fig. 3E. Isoform expression was probed with anti-FLAG antibody, GAPDH served as a loading control. (D) CPT-induced nascent DNA degradation assessed by DNA fiber analysis. SAMHD1-knockout (KO) cells, and SAMHD1-KO cells expressing human SAMHD1 (hSAMHD1), murine SAMHD1 isoform 1 (mSAMHD1 iso1), or isoform 2 (mSAMHD1 iso2) were analyzed following sequential CldU and IdU incubation. Cells were subsequently treated with 1 µM camptothecin (CPT) to induce replication stress, with DMSO-treated cells serving as controls corresponding to Fig. 3F. Nascent DNA degradation was assessed using IdU/CldU tract-length ratios (D) and IdU track-lengths (E) (150 replication forks, two slides for each condition). Statistical significance was determined using the Kruskal–Wallis test followed by Dunn’s multiple-comparisons test (*P < 0.05, ***P < 0.001, ****P < 0.0001). (F) Representative immunoblot corresponding to Fig. 3F and Supplementary Fig. 3D and 3E. Expression of FLAG-tagged hSAMHD1, mSAMHD1 iso1, and mSAMHD1 iso2 in HEK293T SAMHD1-KO cells was detected using an anti-FLAG antibody. GAPDH served as a loading control.

